# Reduced listener-speaker neural coupling underlies speech understanding difficulty in older adults

**DOI:** 10.1101/2020.11.30.403287

**Authors:** Lanfang Liu, Hehui Li, Xiaowei Ding, Qi Zhou, Dingguo Gao, Chunming Lu, Guosheng Ding

## Abstract

An increasing body of studies have highlighted the importance of listener-speaker neural coupling in successful speech communication. How this mechanism may change with normal aging and the association of this change with age-related decline in speech understanding remain unexplored. In this study, we scanned with fMRI a young and an older speaker telling real-life stories, and then played the audio recordings to groups of young (N = 28, aged 19-27y) and older adults (N = 27, aged 58-75y) during scanning, respectively. The older listeners understood the story worse than the young, and the advancing age of the older listeners was associated with poorer speech understanding. Compared to the young listener-speaker dyads, the older dyads exhibited weaker neural couplings in both linguistic and extra-linguistic areas. Moreover, within the older group, the listener’s age was negatively correlated with the overall strength of interbrain coupling, which in turn was associated with poorer speech understanding. These results reveal the deficits of older adults in achieving neural alignment with other brains, which may underlie the age-related decline in speech understanding.

## 1. Introduction

Narratives, especially real-life stories, are the most ancient and effective tools for communicating personal experiences, emotions, and beliefs between people in daily life. To successfully understand spoken narratives requires a set of complex cognitive and neural computations. A listener needs to not only segregate the audio streams into linguistic units and retrieve semantic information, but also simultaneously maintain the decoded information in working memory, integrate it with preceding information and the knowledge base, build situation models, make inferences and predictions, and beyond. Despite these cognitive requirements, young adults can decipher the information intended by a speaker effortlessly. However, the older adults typically understand spoken narratives slower, make more errors, and remember less information (DeDe & Flax, 2016; Schneider et al., 2002; Zacks et al., 2006), even though their basic linguistic functions (such as semantic and syntactic processing) are thought to be preserved (Shafto & Tyler, 2014; Tyler et al., 2010).

Currently, neuroimaging studies on language processing in older adults have primarily focused on the cortical responses to isolated, decontextualized phonemes, words, or sentences within isolated brains (e.g, Anderson et al., 2012; Du et al., 2016; Wong et al., 2009). Little is known about the neural processes underlying naturalistic speech understanding in aging brains. By recording brain activities from both speakers and the listeners engaged in naturalistic verbal communication (i.e., the two-brain approach), recent studies on the young adults have consistently demonstrated that the listeners’ neural activities during speech comprehension were tightly coupled (temporally or spectrally correlated) with the activities in the speaker’s brain during speech production (Dai et al., 2018; Y. Liu et al., 2017; Pan et al., 2020; Silbert et al., 2014; Stephens et al., 2010). Moreover, the tighter listener-speaker neural couplings were associated with better speech understanding (L. Liu et al., 2020; Stephens et al., 2010). The coupled brain activities between communicators are suggested to reflect their alignment at multiple levels of linguistic and extralinguistic representations (Menenti et al., 2012; Schoot et al., 2016). According to psycholinguistic theories and neurocognitive opinions, the alignment between interlocutors can reduce the computational burden involved in language processing, improve neural efficiency and support the building of common conceptual space which lead to mutual understanding (Pickering & Garrod, 2004; A. Stolk et al., 2016; Wheatley et al., 2012).

Up to now, the investigation of human brain in communicative settings has been mainly limited to the young cohorts. How brain-to-brain coupling may change with normal aging and the association of this change with age-related decline in speech understanding remain unexplored. Two possibilities exist. First, due to impaired auditory temporal processing and slowing in cognitive processing, as well decline in high-level cognitive functions (e.g., working memory and attention), an older listener may have difficulties in tracking the fleeing linguistic input and therefore fail to achieve alignment with a speaker on neural activities and mental representations. This failure may in turn lead to poorer speech understanding. Alternatively, according to the interactive brain hypothesis, interactive experience and skills play enabling roles in both the development and current function of social brain mechanisms (Di Paolo & De Jaegher, 2012). In relation to aging, this experience-driven model would predict that the older listeners will exhibit preserved or even tighter neural alignment with the speaker during communication, due to their life-long experience of social interactions through verbal language. The preserved or strengthened ability to achieve alignment with other brains (and minds) may serve as a compensation mechanism to alleviate the adverse effect of cognitive and auditory declines during aging.

To test these possibilities, we scanned a speaker telling real-life stories and later a group of participants listening to the audio recordings of those stories using fMRI. This procedure was applied to the young and older adults separately. Then neural coupling between each listener and the speaker was assessed by calculating the temporal correlation between their BOLD signals in spatially corresponding cortical regions. To determine the potential effect of age on brain-to-brain coupling, we compared the older listener-speaker dyads with the young dyads and further examined the correlation between the age of older listeners and their neural alignments with the speaker. Next, we investigated whether changes in interbrain coupling played a role in age-related decline in speech understanding. To this end, we examined the correlations among age, interbrain coupling, and the level of speech understanding, and then conducted a mediation analysis. If life-long experiences of verbal interactions play a protective role, the older listener-speaker dyads would exhibit comparable or tighter interbrain coupling than did the young dyads. Moreover, the listener-speaker neural coupling would increase with the advancing age of the older listener, which would in turn be associated with better speech comprehension. Alternatively, the older dyads would exhibit weaker neural coupling than did the young, and the age of the older listeners would be negatively correlated with their neural alignments with the speaker. The reduced neural alignment would in turn be linked to poorer speech understanding.

## 2. Method

### 2.1 Participants

Thirty-two young adults (16 females, aged 19-27y) and 30 older adults (18 females, aged 58-75y) participated in this study. Each group included a female speaker, and the rest were listeners. We choose to have the listener and speaker age-matched, rather than letting both young and older adults listen to the same speaker, for two considerations. First, in daily-life situations, verbal communication among older adults differs substantially from that among young adults in terms of topics, speech rate, vocabulary, and organizational structure (Bortfeld et al., 2001; Juncos-Rabadán et al., 2005). Thus, age-matched dyads are likely to communicate better than old-young dyads. Second, the rate of blood flow, metabolism, and neurovascular coupling in the aging brain are quite different from those in younger brains (D’Esposito et al., 2003; Meltzer et al., 2003; Takada et al., 1992); thus, neural dynamics of age-matched brains should be more alike. Given these two points, we presume age-matched listener-speaker dyads are more likely to achieve neural alignment than old-young dyads.

All participants were right-handed based on the Edinburgh Handedness Inventory (Oldfield, 1971), and reported no mental or neurological disorders. For the older group, a Mini-Mental State Examination was administered, and all participants scored above 26. To exclude participants with potential hearing loss, pure tone audiometry was applied. A threshold of < 30 dB HL in the better ear across 0.5, 1.0, 2.0, and 4.0 kHz, which covers the most range of human speech (Turner & Cummings, 1999), was used to filter participants (Peelle & Wingfield, 2016). All participants in the young group met this criterion at every frequency. In the older group, all met this criterion at 0.5, 1, and 2 kHz, and all but two met at 4 kHz. A Digit Span Forward and Backward Memory test was administrated to measure participants’ memory span. Written informed consent was obtained from all participants under the protocol approved by the Reviewer Board of Southwest University. The data of the older group were also used in another study which addressed a different question (L. Liu et al., 2020).

### 2.2 Experimental Design

During the fMRI scanning, the two speakers told stories based on their personal experience, with each story lasting 10 min. To obtain good quality of audio recording, we used a noise-canceling microphone (FOMRI-III, Optoacoustics Ltd., Or-Yehuda, Israel) positioned above the mouth of the speaker, and further de-noised the audio recordings offline using Adobe Audition 3.0 (Adobe Systems Inc., USA). Two groups of college students (N1 = 32, N2 = 31) assessed the complexity and vividness of the stories told by the young and older speakers, respectively, on a scale of 1.0-10. The results showed, different stories told by the same speaker did not differ in either vividness or complexity. Compared to the stories told by the young speaker, the stories told by the older speaker were more vivid (*p* < 0.001) but did not differ in complexity (*p* = 0.25).

Since the older speaker had a sudden cough at the last minute of the recording, we played back only the first 7-min audio recordings to the older listeners during the fMRI scanning. For the young group, the full-length (10 min) recordings were played back. The audio stimuli were presented through an MR-compatible headphone with noise-cancellation (OptoACTIVE, Optoacoustics Ltd., Or-Yehuda, Israel) at a volume comfortably audible for all listeners. A fixation cross lasting for 20s and then an icon of a horn lasting until the end of the scanning were presented to both speakers and listeners (Fig.1). The participants were informed to start speaking or listening immediately upon seeing the horn. To make the participants focus their attention on the speech, they were informed beforehand that an interview about the content of the story would be given after the scanning.

**Fig. 1.**
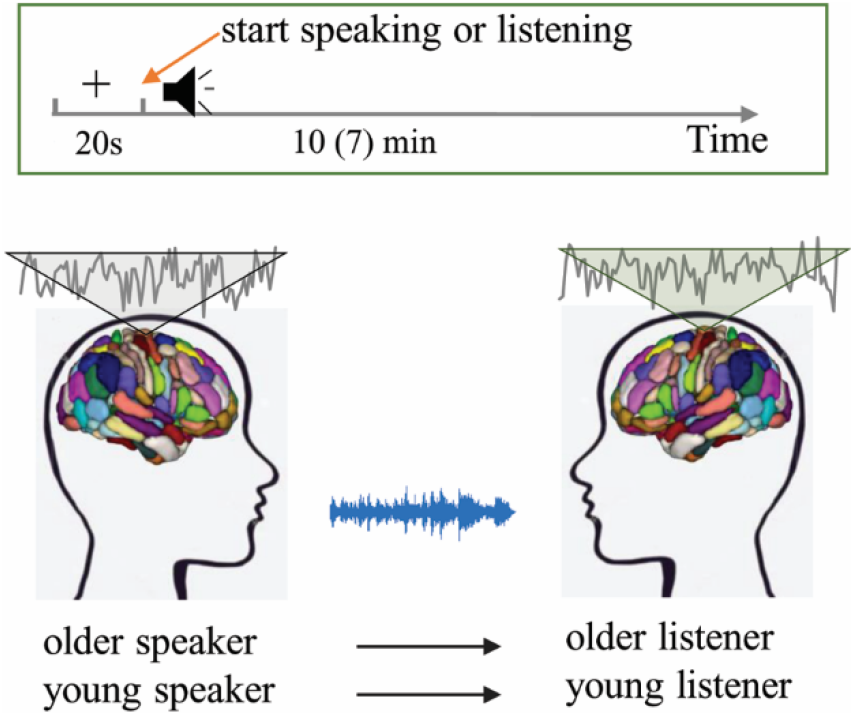
Experimental and analytical protocols. We scanned with fMRI a young and an older speaker telling real-life stories, and then played the recordings to groups of young and older listeners, respectively. In the experiment, a fixation cross lasting for 20s and then an icon of a horn lasting until the end of the scanning were presented to all participants. To assess interbrain coupling, the brain of each participant was first partitioned into 246 regions based on the Brainnetome Atlas, and Pearson’s correlations were calculated between the mean signals of each region from the listener’s brain and those of the homologous region from the speaker’s brain.

Each young adult listened to one of the three stories told by the young speaker. Each older adult listened to one of the two stories told by the older speaker. As participants listening to the different stories did not differ in their comprehension score (for young group: *F*_(2, 29)_ = 2.96, *p* = 0.07; for older group: *t*_(29)_ = −0.18, *p* = 0.85), their data were collapsed for further analyses.

### 2.3 Behavioral assessment for the level of speech understanding

Two measurements were used to assess the level of speech understanding. First, at the end of play, listeners in the scanner were asked to report to which degree (in the form of percentage) they understood the story. To obtain a more objective assessment, the listeners were required to retell the stories in detail immediately after the scanning. After the free recall, the experimenters asked the participants several questions regarding the part of the content not present in the recall. Two independent raters then scored each listener based on their free call and their answers to the questions. Details for the scoring procedure were described in the supplementary material. As the assessments made by the two raters were in high agreement (Spearman’s r_(62)_= 0.85), we averaged the two scores to quantify the listener’s level of speech understanding.

### 2.4 MRI acquisition and preprocessing

Imaging data were acquired with a 3T Siemens Trio scanner in the MRI Center of the Southwest University of China. A gradient echo planar imaging sequence was applied to collect functional images. The parameters were: repetition time = 2000 ms, echo time = 30 ms, flip angle = 90°, field of view = 220 mm, matrix size = 64 × 64, slice number = 32 interleaved, voxel size = 3.44 × 3.44 × 3.99 mm^3^. A MPRAGE sequence was adopted to collect T1 structural images with the following parameters: repetition time = 2530 ms, echo time = 3.39 ms, flip angle = 7°, FOV = 256 mm^2^, scan order = interleaved, matrix size = 256 × 256, and voxel size = 1.0 × 1.0 × 1.33 mm^3^.

A total of 310 and 220 volumes were acquired for the young and older participants, respectively. The first ten volumes corresponding to the fixation period were discarded. Image preprocessing was implemented using the DPABI toolkit (Yan et al., 2016) which is based on SPM12 (www.fil.ion.ucl.ac.uk/spm/) modules. First, slice-timing correction was performed to correct for varied sampling time of slices. Next, the corrected images were spatially realigned and co-registered to individual subjects’ anatomical images. The resultant images were then spatially normalized to Montreal Neurological Institute (MNI) space, resampled into a 3 × 3 × 3 mm^3^ voxel size, and spatially smoothed using a 7 mm full-width-half-maximum (FWHM) Gaussian kernel. Finally, the preprocessed images were detrended, nuisance variable regressed, and high-pass filtered (1/128 Hz). The nuisance variables included five principal components of white matter and cerebrospinal fluid within individual subjects’ T1 segmentation mask (Behzadi et al., 2007), as well as Friston’s-24 motion parameters (Friston et al., 1996). The datasets of four young and three older participants were discarded due to excessive head movement (more than 3mm or 3 degrees).

### 2.5 Data analysis

#### 2.5.1 Analysis of behavioral data

To test whether speech understanding declined with aging, we compared the older adults with young on the self-reports and comprehension scores using two-sample *t*-tests. In addition, we examined whether the increased age of the older adults was associated with a lower comprehension score. As participants’ memory capacity may influence their recall for the speech contents, here we applied a Spearman’s partial correlation which included the digit spans on the forward and backward memory test as two covariates. In the partial correlation analysis, the digit spans were regressed out separately from age and comprehension scores, and the residuals of the two models were used to compute the correlation value. Since the directions of between-group differences and the brain-behavior correlation had been predicted, a one-tailed test for statistical significance was employed.

#### 2.5.2 Measurement for listener-speaker neural coupling

The listener-speaker neural coupling was measured by the Pearson’s correlation coefficients between the time series of each area in the listener’s brain and the time series of the homologous area in the speaker’s brain (Fig 1). Since there are noticeable intersubject variability in brain functioning and structural morphometry among older adults (Geerligs et al., 2017; Kannurpatti et al., 2010; Raz et al., 2010), functionally similar loci may not correspond well in spatial location across older adults. To alleviate the potential influence of inter-individual variability, we computed the interbrain correlation at the regional level, which is the common practice in studies on intra-brain functional connectivity. For each participant, the brain was partitioned into 246 regions based on the Human Brainnetome Atlas (Fan et al., 2016). At each region, the time series across all voxels within it were averaged for calculating the interbrain correlation. To test the robustness of the results, we re-analyzed the data using Craddock’s atlas (Craddock et al., 2012) which partitioned the brain into 200 regions and obtained quite similar results (supplementary material).

As listeners may need time to extract high-level information in the speech, their neural activities would lag those in the speaker’s brain. Besides, listeners may anticipate the speech content, their neural activities would precede those in the speaker’s brain. To capture the temporal asymmetry in neural coupling, we repeated the interbrain correlation analysis by shifting the listener’s time course with respect to those of the speaker from −8s (listener preceding) to 8s (listener lagging) in 2s increments. At shift zero, the listener’s brain activity was time-locked to the speaker’s vocalization. To obtain an overall temporal profile of the interbrain coupling and identify the time point wherein this effect reached the peak (referred to as the “peaking time”, which was the target of the bellowing analysis), we calculated the mean of the correlation values across the 246 parcels and all participants at each lag.

At each lag, a one-tailed *t*-test was performed on the interbrain correlation coefficients (the r values) for the young and older groups separately. The resulting coupling maps were corrected for multiple comparisons using the Benjamini-Hochberg procedure to control the false discovery rate (FDR) at 0.05.

#### 2.5.3 Evaluation for the effect of age on listener-speaker neural coupling

Two folds of analyses were conducted to assess the effect of age on listener-speaker neural coupling. First, we directly compared the older group to the young group at each time lag using a two-sample *t*-test. Nevertheless, the between-group effects we may find could be driven by not only the differences in the listener’s age but also the differences in the speaker and the speech contents. To eliminate the confounding effect, we further assessed the correlation between the chronological age of older listeners and their neural alignment to the speaker. To measure the overall strength of neural alignment, we selected the collection of cortical regions showing the most significant listener-speaker neural coupling (for the peaking time) at the group level. Then for each participant, the mean of the correlation values within these regions was calculated. Before averaging, each r-value was transformed to z-values using Fisher’s z transformation to normalize its distribution. The averaged z values were then inverse transformed (z-to-r) to produce average r values. To ensure that the number of selected cortical regions did not have an undue influence on the results, we repeated this analysis over a wide range of threshold: from the top 10 regions showing the strongest interbrain couplings (corresponding *t*_(26)_ > = 2.85) to the top 44 regions (corresponding to *t*_(26)_ > =1.65). The MNI coordinates and anatomical labels for the 44 regions were reported in the supplementary material (table S1).

#### 2.5.4 Brain-behavior correlation

To assess the association of listener-speaker coupling with the listener’s level of speech understanding, we computed the Spearman’s rank correlation between the comprehension scores and the overall strength of interbrain coupling. To control for the potential effect of memory capacity on the behavioral measurement for speech understanding, a partial correlation was adopted which took digit spans on the forward and backward memory test as two covariates. Consistent as above, we took the mean value from the collection of cortical regions demonstrating the most significant interbrain coupling and repeated the correlation analyses over a wide range of thresholds.

#### 2.5.5 Mediation analysis

To test whether the interbrain coupling mediated the relationship between the older listener’s age and the level of speech understanding, we performed a mediation analysis using the Hayes’ PROCESS macro in SPSS. In the analysis, the age of the older listeners and their comprehension scores were entered as the causal variable and the outcome, respectively, and the overall strength of listener-speaker neural coupling was entered as the mediating variable. A series of linear regression were used to assess (1) the effect of age on interbrain coupling (denoted as *a*); (2) the effect of interbrain coupling on speech understanding after removing the effect of age (denoted as *b*); and (3) the total effect of age on comprehension score (denoted as *c*). The mediation effect of interbrain neural coupling on the age-behavior relationship was assessed by taking the product of coefficient *a* and coefficient *b* (denoted as *ab, also called “indirect effect”*). In all three regression models, the backward and forward memory spans of older listeners were included as two covariates. A bootstrapping procedure with 5,000 iterations was used to assess the 95% confidence interval for *ab* (Hayes & Scharkow, 2013). The indirect (mediation) effect was statistically significant if the confidence interval did not include zero. For the mediation analyses, we examined and reported the collection of brain regions (the top 34) showing the strongest positive correlation with the comprehension score. We also examined other collections and obtained similar results.

### 2.6 Analyses for potential factors contributing to the aging effect on interbrain coupling

We explored three factors that potentially account for the relatively weak neural coupling between the older listener-speaker dyads, including poor sustained attention, reduced cortical responsiveness to the speech, and age-related gray matter atrophy. Details for the following analyses were provided in the supplementary material.

#### 2.6.1 Potential contribution of poor sustained attention

It has been well-documented that older adults have difficulties in sustaining their attention over time (McAvinue et al., 2012; Staub et al., 2014). It is possible that the older listener-speaker dyads presented as tight interbrain coupling as the young dyads at the beginning of the communication, which dropped down gradually due to that the older listeners could not well sustain their attention on the speech. In the above analyses, the interbrain coupling was assessed using the full-length time series, thus reflecting a relatively average status over the verbal communication. To investigate the potential role of sustained attention, we performed a sliding window analysis to capture the time-dependent variations in the interbrain effect.

#### 2.6.2 Potential contribution of age-related reduction in cortical responsiveness to the speech

It has been reported that several cortical regions were less activated in older adults than in the young during language processing (Manan et al., 2015; Wong et al., 2009; Zlatar et al., 2013). Reduced cortical responsiveness to the speech signal in the listener’s brain may lead to the weak listener-speaker neural coupling. To test this possibility, we conducted inter-subject correlation (ISC) analyses to identify brain regions that process the stimulus in a consistent, time-locks manner (Nastase et al., 2019), and examined for each brain region the correlation between ISC value and listener-speaker coupling strength across participants.

#### 2.6.3 Potential contribution of age-related gray matter atrophy

Finally, we explored whether the deficits of aged brain in “tuning into” other brain was associated with age-related gray matter atrophy. We first tested whether those cortical regions showing age-related decline in neural coupling also exhibited age-related reduction in gray matter volume (GMV). We then examined for each cortical region whether the strength of interbrain coupling was correlated with the GMV of this region across the participants.

## 3. Results

### 3.1 Behavioral results

Despite the stories told by the older speaker were more vivid and shorter than those told by the young speaker, whereas the complexity of speech content did not differ between the two, the older listeners still understood the speech worse than the young, as demonstrated in both the self-reports (two-sample *t*_(47)_ = 5.95, p < 10^−6^) and comprehension scores (two-sample *t*_(53)_ = 5.80, *p* <10^−6^). Within the older group, the comprehension score was significantly correlated with the listener’s backward memory span (r_(25)_ = 0.50, *p* < 0.01, by Spearman’s correlation), but not with the forward memory span (r_(25)_ = 0.133, *p* > 0.5) or hearing level (r_(25)_ = −0.12, *p* > 0.5). After controlling for the individual differences in memory spans, the increasing age of the older listener was associated with a lower comprehension score (Spearman’s partial r_(25)_ = −0.37, one-tailed *p* = 0.035) (Fig.2). These results are consistent with previous reports that the ability to comprehend spoken language declines with aging.

**Fig. 2.**
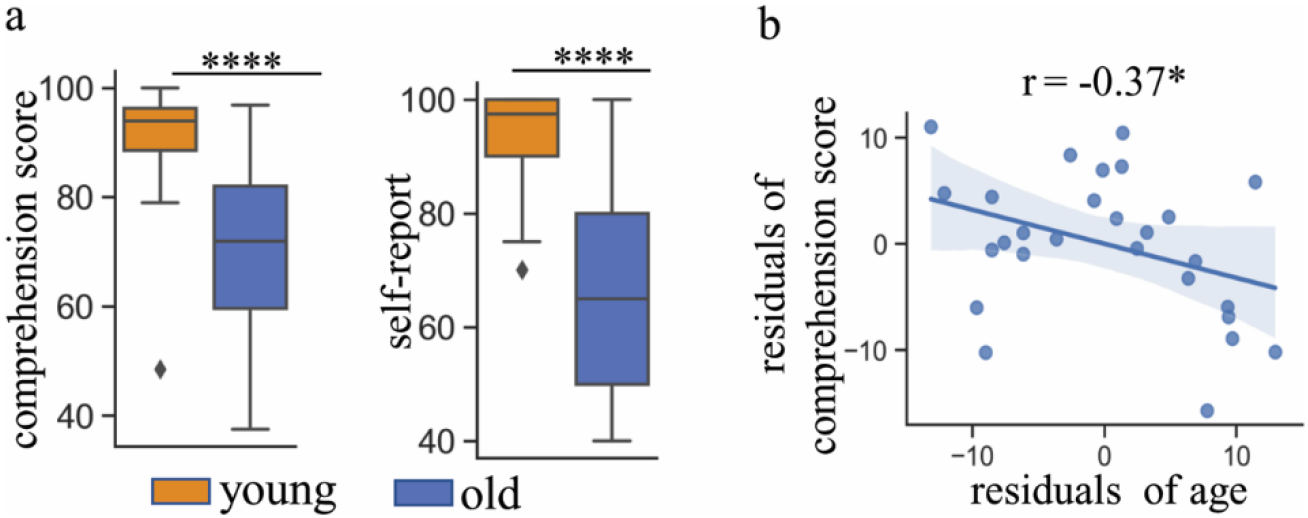
Behavioral results. (a). Older adults understood the narratives poorer than did the young, as demonstrated by both self-reports and comprehension scores. (b). The age of the older listener was negatively correlated with the comprehension score, after controlling for the effect of memory spans. *: *p* < 0.05; ****: *p*< 0.0005.

### 3.2 Widespread neural coupling between the young listener and speaker

Consistent with prior work (Stephens et al., 2010; Arjen Stolk et al., 2014), we found widespread neural couplings between the listener and speaker from the young group. On average, the activity in the listener’s brain was aligned to the activity in the speaker’s brain with some temporal delays. The strongest neural coupling at the whole-brain level occurred at a time lag of 6s (Fig.3). Notably, the listener’s neural alignment with the speaker followed an orderly temporospatial progression: shortly after the speaker’s vocalization (0-2s), activities in the lower-order auditory areas (the superior temporal gyrus and sulcus, STG/STS) of the listener’s brain were aligned to the activities of the homologous regions in the speaker’s brain; 4-6s latter, the higher-order semantic/conceptual areas, encompassing bilateral inferior parietal lobule (IPL), inferior frontal gyrus (IFG), angular, bilateral precuneus, posterior cingulate gyrus (PCG) were aligned (Fig. 3). For delays longer than 6s, the interbrain coupling declined substantially. When the listener’s brain activities were shifted to precede those in the speaker’s brain, no significant interbrain correlation was found. Thus, further analyses only focused on the interbrain coupling with a lag from 0s to 6s.

**Fig. 3.**
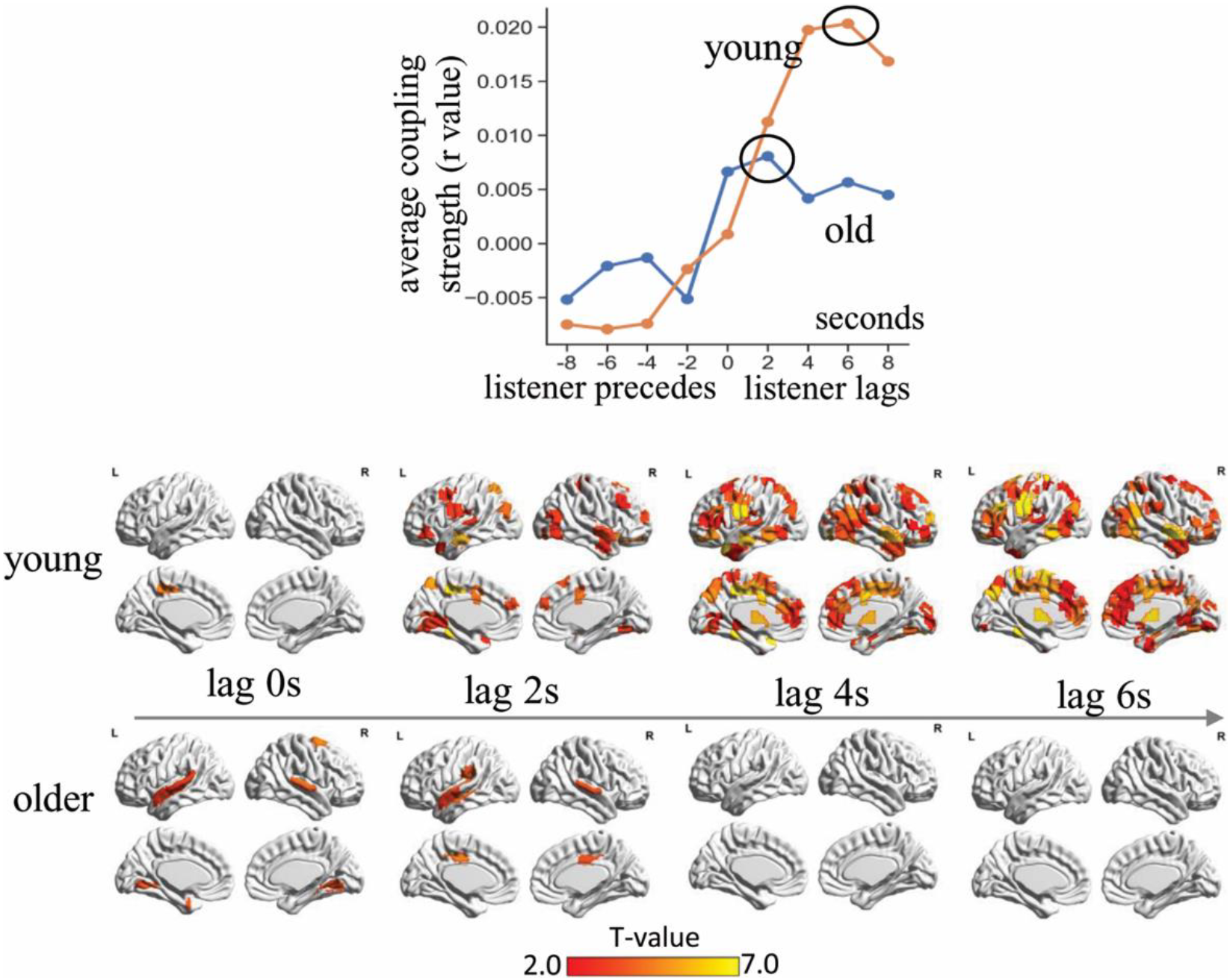
The temporospatial pattern of listener-speaker neural coupling. In both groups, the listener’s brain activities got aligned with the speaker’s brain activities with some temporal lags. Compared to the young listener-speaker dyads, neural couplings between the older dyads were less extensive and peaked earlier (6s versus 2s). The upper panel shows the coupling strength averaged across the whole brain and across all subjects. The lower panel displays the *t*-statistic maps thresholded with FDR corrected *p* < 0.05.

### 3.3 Weak neural coupling between the older listener and speaker

In contrast to widespread interbrain couplings in the young group, the older listener-speaker dyads showed significant neural couplings in only a few cortical regions, including the bilateral STG, PCG, lingual gyrus, and the left supramarginal gyrus. As with the young group, in the older group, the listener’s brain activity got aligned with the activities in the speaker’s brain with some temporal lags (Fig. 3) but peaked earlier (at a lag of 2s) and faded quicker.

### 3.4 listener-speaker neural coupling decreased with advancing age

The two-sample *t*-test revealed that neural couplings between the older dyads were significantly weaker than the young dyads in multiple cortical regions, including the bilateral STG/STS, superior and middle frontal gyrus, angular gyrus, and precuneus. In line with the results of between-group comparisons, the age of the older listeners was negatively correlated with the overall strength of interbrain coupling. This effect was robust to the changes of selected cortical regions showing significant interbrain couplings (Fig. 4).

**Fig. 4.**
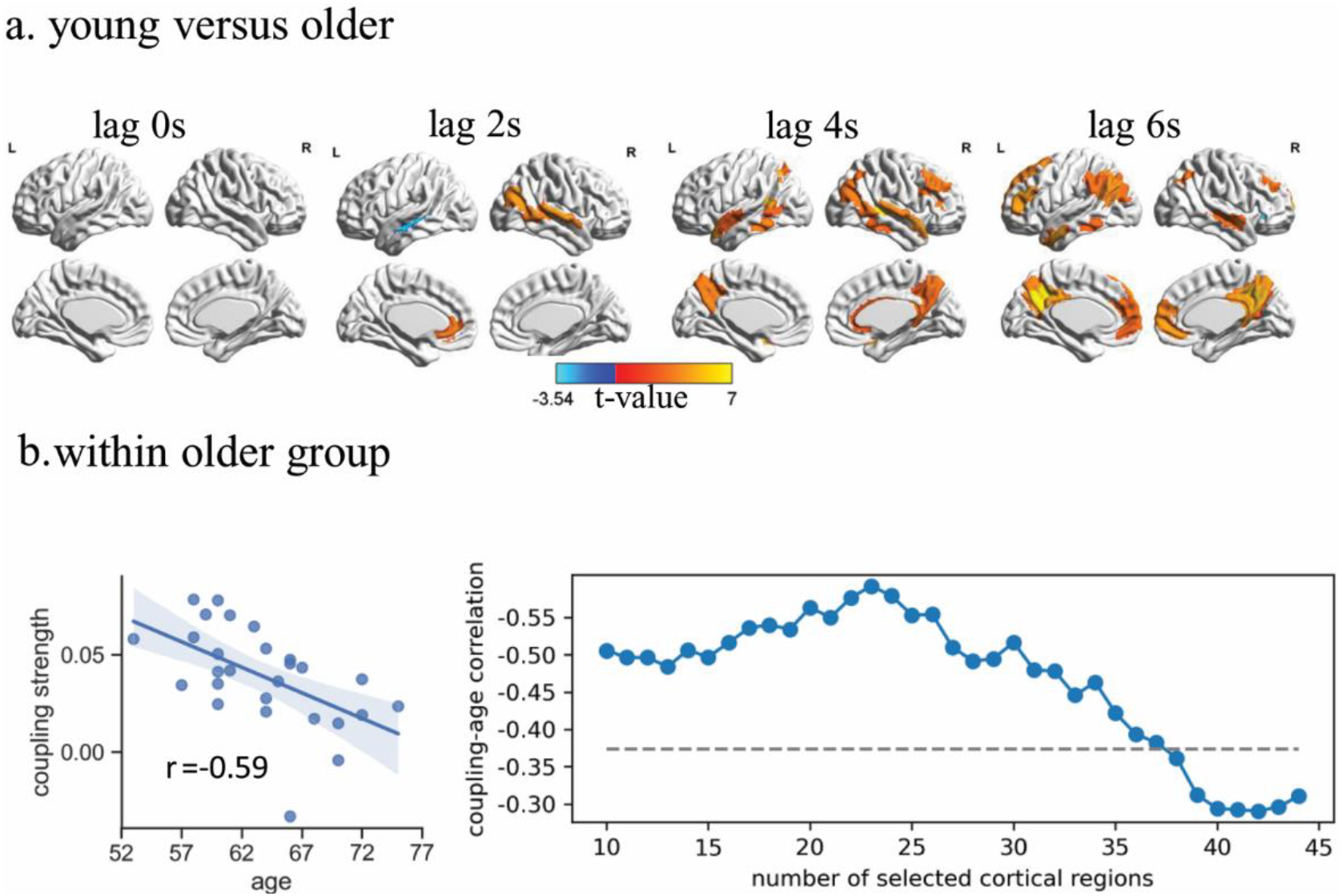
Reduced listener-speaker neural couplings with aging. (a). Direct contrast between the young and the older group. Orange: young > older; Blue: older > young. Threshold: FDR corrected *p* < 0.05. (b). The overall strength of interbrain coupling declined as a function of the older listener’s age. *Left*: scatter plot showing the negative correlation between the older listener’s age and the overall strength of listener-speaker neural coupling. *Right*: This effect was consistent across a wide range of selected cortical regions. The vertical line indicates the r value corresponding to *p* = 0.05.

### 3.5 Correlation between neural coupling and speech understanding

Consistent with previous studies, we found the overall strength of listener-speaker neural coupling was positively correlated with listeners’ comprehension scores (with memory span controlled) (Fig.5). This relationship was found in both the young and older groups and was consistent across a wide range of selected cortical regions. In the young group, it was the 6s-lagged neural coupling (the peaking time) that demonstrated the relevance to speech understanding. In the older group, it was the 2s-lagged (the peaking time) neural coupling that correlated to speech understanding.

**Fig. 5.**
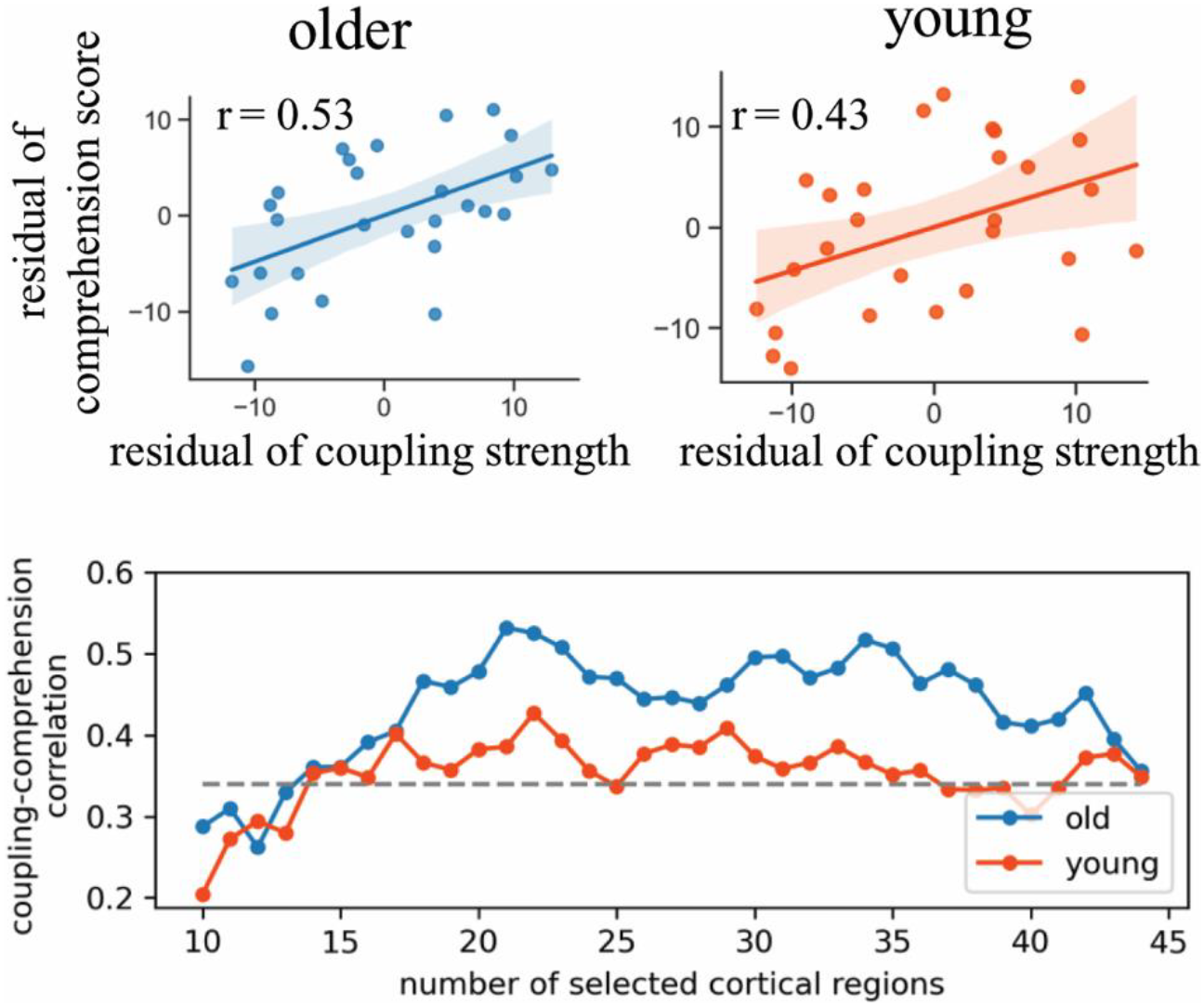
Stronger listener-speaker neural coupling was associated with better speech understanding in both the young and older groups. *Upper:* Scatter plots showing a significant positive correlation between the overall strength of listener-speaker neural coupling and the listener’s comprehension score, with the effect of memory spans being controlled. *Lower:* This effect was consistent across a wide range of selected cortical regions. The vertical line indicates the r value corresponding to *p* = 0.05.

### 3.6 Reduced interbrain coupling mediated the negative relationship between age and speech understanding

In the above analyses, we observed in the older group 1) a negative correlation between age and speech understanding; 2) a negative correlation between age and interbrain coupling; and 3) a positive correlation between interbrain coupling and speech understanding. Based on these findings, it was plausible to assume a mediation role of interbrain coupling in the relationship between age and speech understanding. Indeed, the mediation analyses revealed a significant mediating effect of interbrain coupling on the relationship between age and speech understanding (*ab* = −0.62, 95% confidence interval: [-1.32 to −0.05], obtained from the Bootstrapping test) (Fig. 6). Moreover, after controlling for the effect of neural coupling, the listener’s age was no longer a significant predictor for the listener’s comprehension score. These results indicate the advancing age of the older listener was associated with poorer ability to achieve neural alignment with the speaker, which in turn negatively affected the listener’s speech understanding.

**Fig. 6.**
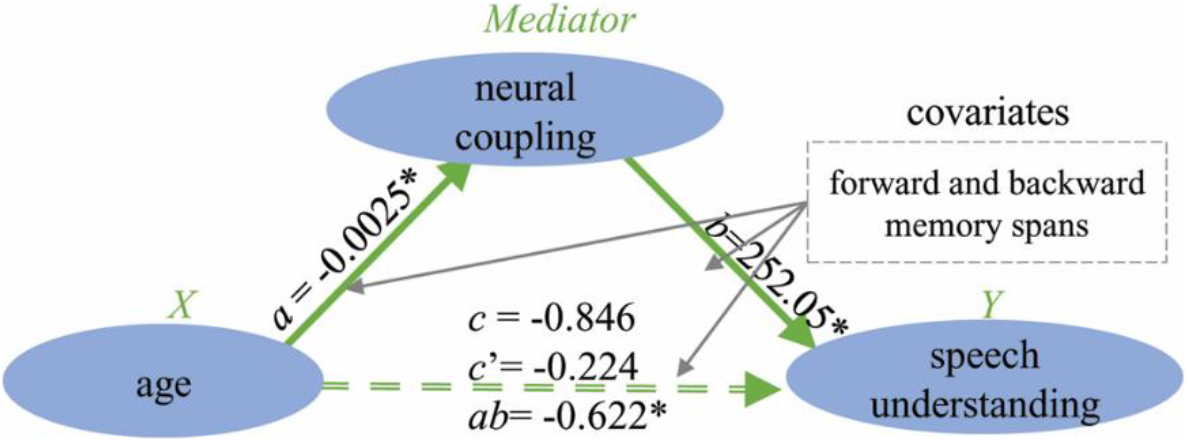
Reduced interbrain coupling mediated the negative relationship between age and speech understanding. *a*: the negative effect of older listener’s age on listener-speaker neural coupling. *b*: the positive effect of neural coupling on speech understanding after controlling for the effect of age. *c*. The total effect of age on speech understanding. *c*’: the direct effect of age on speech understanding. *ab*: the indirect effect of age on speech understanding through interbrain coupling.

### 3.7 Potential factors contributing to the aging effect on interbrain coupling

#### 3.7.1 Neural coupling between the older dyads decreased over time

The sliding-window analyses revealed, the older listener-speaker dyads presented extensive neural couplings within the first 2 min of the communication, which gradually decreased thereafter. In comparison, the young dyads exhibited extensive neural couplings across multiple time windows (Fig. S1), and the extent of interbrain coupling tended to increase over the course of the communication. These results indicate that the poor ability to keep attention to the speech may partially account for the weak neural alignment of the older listener with the speaker.

#### 3.7.2 Widespread listener-listener neural correlations in the older group

`The ISC analyses for the older group revealed, the neural activities in the listener’s brain were correlated significantly with the brain activities of other participants listening to the same story. Significant listener-listener neural correlations were found in both linguistic and multiple extra-linguistic areas, suggesting active involvement of those regions in speech processing (Fig. S2). Throughout the whole brain, there was no region showing significant correlation (FDR corrected *p* < 0.05) between the strength of listener-speaker neural coupling and the strength of listener-listener neural correlations. These results suggest the weak neural alignment of the older listener with the speaker was unlikely to arise from age-related reduction in cortical response to the speech.

#### 3.7.3 Reduced interbrain coupling was accompanied by gray matter loss

On those cortical regions of which the listener-speaker neural coupling decreased with age, we also observed a significant negative correlation between the listener’s age and the average GMV. This means, the same network showing age-related decline in interbrain coupling also exhibited age-related gray matter loss. Still, this negative relationship between age and GMV was consistent across a wide range of selected cortical regions (Fig. S3). Nevertheless, we did not find a linear correlation for any of the 246 brain regions between the strength of interbrain coupling and the GMV across the participants, indicating a complicated relationship between changes in brain function and brain structure during normal aging.

### 3.8 Validating the main findings with another brain parcellation scheme

We repeated the above analyses using the human brain atlas proposed by Craddock *et.al* which partitioned the brain into 200 regions (Craddock et al., 2012). The results of interbrain coupling (Fig.S4), age-coupling correlation (Fig.S5), comprehension-coupling correlation (Fig. S6), and mediation analyses (Fig.S7) were consistent with those using the Human Brainnetome Atlas.

## 4. Discussion

This study investigated how neural coupling between the listener and speaker during verbal communication, an important mechanism underlying information transfer across brains, may change with normal aging, and the association of this change with age-related decline in speech understanding. We found, compared to the young listener-speaker dyads, verbal communication between the older dyads were less successful, and at the same time, neural couplings between the older dyads were weaker and peaked earlier. Moreover, within the older group, the increasing age of the listeners was associated with weaker listener-speaker neural coupling, in turn associated with poor speech understanding. The implications of those findings are discussed below.

### 4.1 Age-related reduction in listener-speaker neural coupling

Consistent with previous reports (Silbert et al., 2014; Stephens et al., 2010), we observed widespread neural couplings between the young listener-speaker dyads during the communication. However, applying similar experimental protocol and analytical methods, we found quite weak neural couplings between the older listener-speaker dyads, which occurred mainly in the bilateral STG and PCC. Moreover, the overall strength of interbrain coupling was negatively correlated with the age of the older listener. Compared to the young dyads, the older dyads exhibited reduced neural couplings in both frontal-temporal language areas and a set of extra-linguistic areas, including the precuneus, medial prefrontal cortex, and the temporal-parietal conjunction. Those extra-linguistic regions are known to be involved in social cognition critical for interpersonal communication, including mentalizing, simulation, and making inference for other’s emotions, beliefs, goals, and intentions (Spreng & Grady, 2009). The reduced neural couplings indicate the older listeners were less able to achieve alignment with the speaker on either linguistic or higher-level conceptual representations (such as the situation model).

In contrast to the weak listener-speaker neural couplings, we observed strong and widespread listener-listener neural correlations in the old group. Given that those brain regions did not respond to the story were unlikely to fluctuate synchronously across participants, the extensive ISC suggests multiple brain regions in the older adults had been actively recruited for processing the speech. These results also imply that, despite preserving high responsiveness to external stimuli, the aged brains can hardly resonate with and be shaped by other brains during social interactions. Nevertheless, it should be noted that the current study only investigated the communication between older adults. It remains to be seen whether it is also difficult for the older listener to achieve neural alignment with a young speaker. A related question is whether an older speaker was less able to induce resonance and shape the listener’s brain than did the young speaker.

### 4.2 Possible causes for reduced neural alignment in older adults

The reduced neural coupling between the older listener-speaker dyads may arise from age-related declines in peripheral hearing, high-level cognitive functions, as well as regional brain atrophy. It is argued that the brain-to-brain neural couplings are built on the brain-to-stimuli couplings (Hasson et al., 2012), which has been supported by a recent empirical study (Perez et al., 2017). Given that normal aging is accompanied by reductions in neural entrainments to the temporal structure of speech (Henry et al., 2017), it is possible that reduced brain-to-stimuli couplings in the listener (and may also in the speaker) partially lead to the reduced brain-to-brain couplings. Another possible cause is the poor ability of older listeners to keep attention to the external speech. Indeed, the sliding window analyses revealed the extent of neural coupling between the older listener-speaker dyads tended to decrease over the course of communication, while that between the young dyads tended to increase. Finally, age-related brain structural changes may play a role in the reduced interbrain coupling. We found the same network showing age-related decline in interbrain coupling also exhibited age-related reduction in GMV. Nevertheless, we did not find a significant linear relationship between the GMV and interbrain coupling across participants, possibly because there exist complicated nonlinear interactions between brain function and structure during the aging process.

### 4.3 How reduced neural alignment relates to poorer speech understanding

In both the young and older groups, stronger neural alignment of the listener with the speaker was associated with better speech understanding (after controlling for the individual differences in memory span). Currently, despite brain-to-brain couplings have been frequently reported, how this interbrain effect contributes to interpersonal communication remains elusive. Notably, memory research has repeatedly shown that greater neural similarity between encoding and retrieval of an event was associated with better memory for the event (Ritchey et al., 2013; Xue et al., 2010). In addition, it has been demonstrated that neural patterns observed as participants watched a movie were significantly correlated with neural patterns of naïve participants listening to the spoken description of the movie, and greater viewing-listening pattern similarity was associated with listener’s better memory for the speech (Zadbood et al., 2017). Considering the interbrain neural alignment reflects the degree of neural similarity (in the temporal space), an intriguing possibility is that resembling the neural activities in the speaker’s brain can help the listener to store the fleeting speech information into memory. In this study, the weak neural alignments with the speaker may reduce the efficiency of the older listeners to store the currently decoded information into memory, which may lead to less successful online comprehension and subsequently poor recall for the speech contents.

### 4.4 Conclusions and outlook

In summary, this study revealed for the first that the older listeners have difficulties in achieving neural alignment with the speaker during verbal communication, which in turn linked to their poor performance in comprehending and memorizing the speech contents. These findings provide novel insights into the neural basis underlying age-related decline in speech understanding, and suggest the between-brain neural coupling can potentially be used as a biomarker for quantifying the degree to which one understands other people via speech. Future studies should combine the analyses for between-brain interaction and those for within-brain activities to obtain a more complete picture of the neural mechanism underlying speech comprehension and the effect of age on this function. Besides, it would be interesting for future studies to examine the neural coupling between children, together with our findings on the young and older adults, to uncover the developmental trajectory of between-brain interactions.

## Acknowledgments

This work was supported by grants from the National Natural Science Foundation of China (NSFC: 3190082, 31971036), China Postdoctoral Science Foundation (2019M653248) and the Open Research Fund of the State Key Laboratory of Cognitive Neuroscience and Learning of Beijing Normal University (CNLYB1803).

## Credit Author Statement

Guosheng Ding: Conceptualization, Writing-Review & Editing, Supervision; Lanfang Liu: Conceptualization, Formal analysis, Writing-Original Draft, Project administration; Xiaowei Ding: Writing - Review & Editing; Hehui Li: Writing-Review & Editing; Qi Zhou: Project administration; Dingguo Gao: Writing-Review & Editing; Chunming Lu: Conceptualization, Writing - Review & Editing.

