## supplementary material for "Reduced listener-speaker neural coupling underlies speech understanding difficulty in older adults"

### 1. The scoring procedure

To make the scoring as objective as possible, we made a list of questions regarding important contents of the story (such as “what happened during her journey to the airport”). The correct answers included several critical points covering characters, places, time, and motivations et al. A score was given according to the information a listener provided about those key points for each question on the list.

### 2. Analysis of the potential cause of age-related reduction in interbrain coupling

#### 2.1 Sliding window analyses

We performed the sliding window analyses to test the potential impact of sustained attention on listener-speaker neural coupling. In the analysis, the time series of brain activity in each region were segmented into short windows with a length of 30 TR and shifted with a step size of 3 TR, resulting in 59 and 89 windows for the older and young groups, respectively. For each window, Pearson’s correlation between the listener’s and the speaker’s brain activities were calculated using the 30 time points, and one-tailed *t*-tests were performed to identify those cortical regions showing significantly higher interbrain correlation value than zero. Since the purpose of this analysis was to gain a qualitative picture of the dynamic properties of the interbrain effect, here a lenient threshold (p < 0.05, uncorrected) was applied. The extent of listener-speaker neural coupling was determined by the total number of significant cortical regions under each time window. For this analysis, only the brain activities where the interbrain coupling reached the peak (with a 6s-lag for the young and a 2s-lag for the older) were examined.

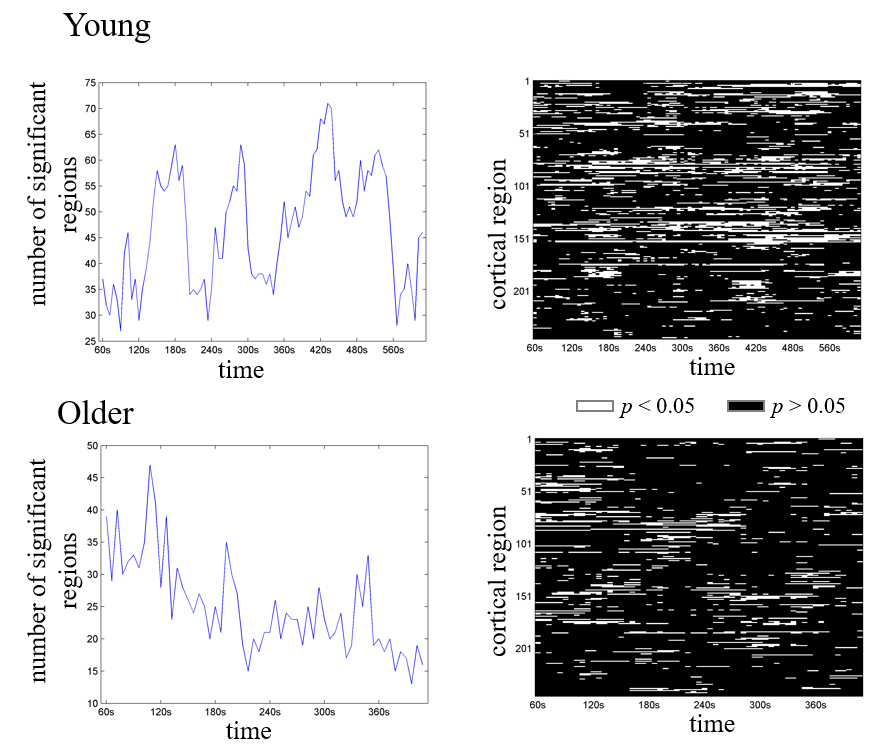

Fig.S1: The dynamics of listener-speaker neural coupling during the communication. In the young group, the listener-speaker dyads consistently exhibited extensive neural coupling over time. However, in the old group, the extent of interbrain coupling tended to decrease over time.

#### 2.2 Listener-listener neural correlation

We conducted inter-subject correlation (ISC) analyses for the older group to identify brain regions that were actively involved in speech processing. For each listener, Pearson’s correlation between the regional time courses and the average time course from other listeners exposed to the same story was calculated. Then a two-tailed *t*-test was conducted on the r values to obtain the ISC map. The results were corrected for multiple comparisons using FDR correction.

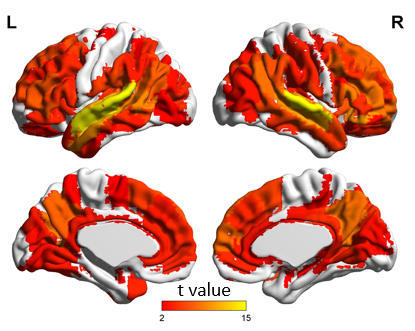

Fig.S2 Extensive neural correlations among the older listeners.

#### 2.3 Analyses for gray matter volume

We explored whether the deficits of older listener in achieving neural alignment with the speaker was related to age-related gray matter atrophy. Gray matter volume was measured via Voxel-based Morphometry (VBM) analysis implemented by the Computational Anatomy Toolbox 12 (CAT12; http://dbm.neuro.uni-jena.de/cat.html). The structural images were preprocessed using the default parameters of this toolbox, including corrections for bias-field inhomogeneities, segmentation into gray matter, white matter, and cerebrospinal fluid, and spatial normalization to the DARTEL template in MNI space. The resulting gray-matter images were modulated in order to preserve the total amount of gray matter signal in the normalized partitions. Careful inspection of quality parameters generated automatically by the toolbox suggests no exclusion of subjects’ VBM data. Consistent with the above analysis, we obtained the representative GMV for each region from the Craddock's atlas by averaging the GMV values over all voxels within the region.

We first assessed the correlation between GMV and age. Consistent with the main analysis, we calculated the mean GMV for the collection of cortical regions showing the most significant interbrain coupling. Next, we analyzed for each region the correlation between the strength of interbrain coupling and the GMV of this region across the listeners. The Pearson’s partial correlation analysis was used here, which included participants’ total intracranial volume obtained from the VBM analysis as a covariate to correct for differences in brain size.

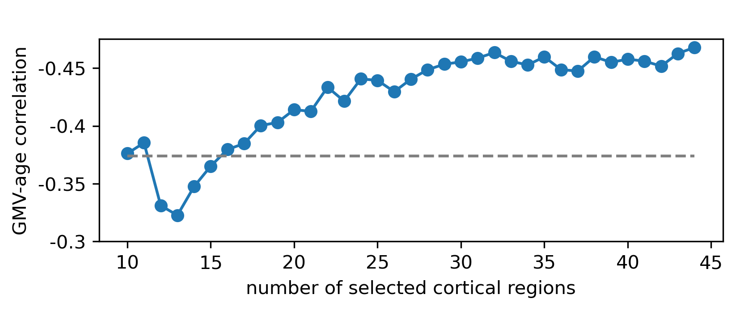

Fig. S3. The same network showing age-related reduction in interbrain coupling also exhibited age-related gray matter loss.

### 3. Validating the main findings with another brain parcellation scheme

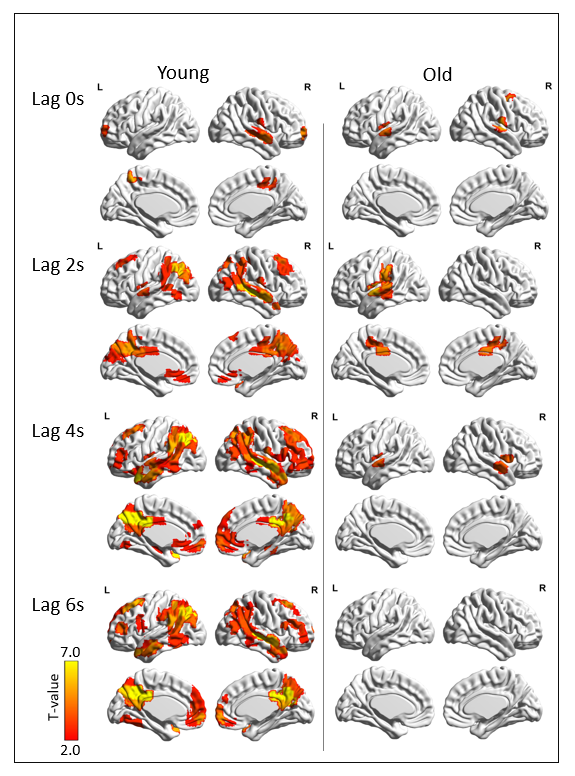

Fig.S4. Listener-speaker neural coupling. Threshold: FDR corrected *p* <0.05.

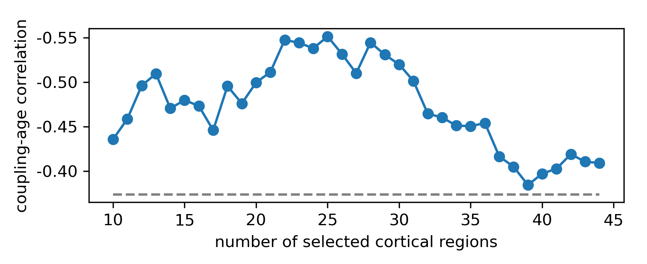

Fig. S5. Negative correlation between the age of older listener and the overall strength of interbrain coupling. This relationship was consistent across a range of selected cortical regions.

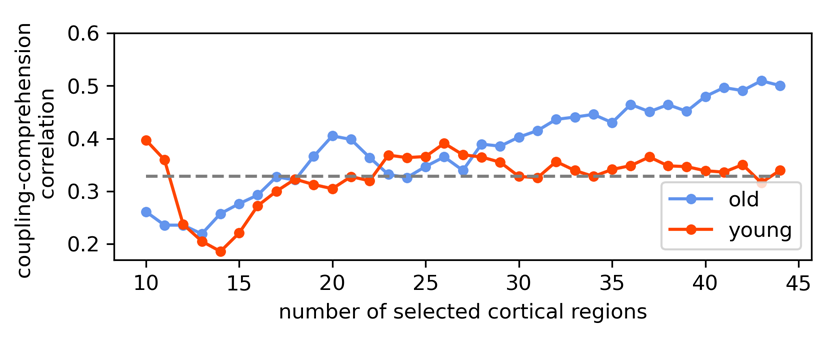

Fig. S6. Positive correlation between comprehension score and the overall strength of interbrain coupling. This relationship was consistent across a range of selected cortical regions.

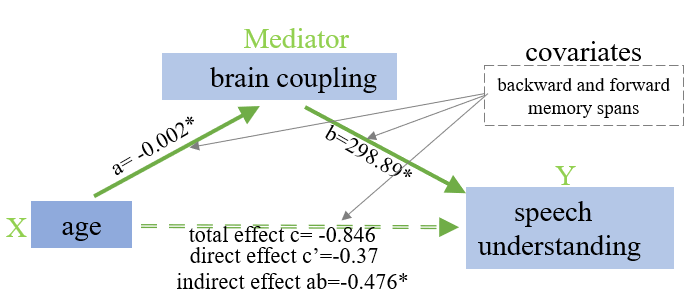

Fig. S7: The mediating effect of interbrain coupling on the relationship between age and comprehension score. The 90% confidence interval for the indirect (mediating) effect was from -1.08 to -0.40.

Table S1: The top 44 regions showing the most significant listener-speaker neural couplings in the young (left) and older (right) groups.

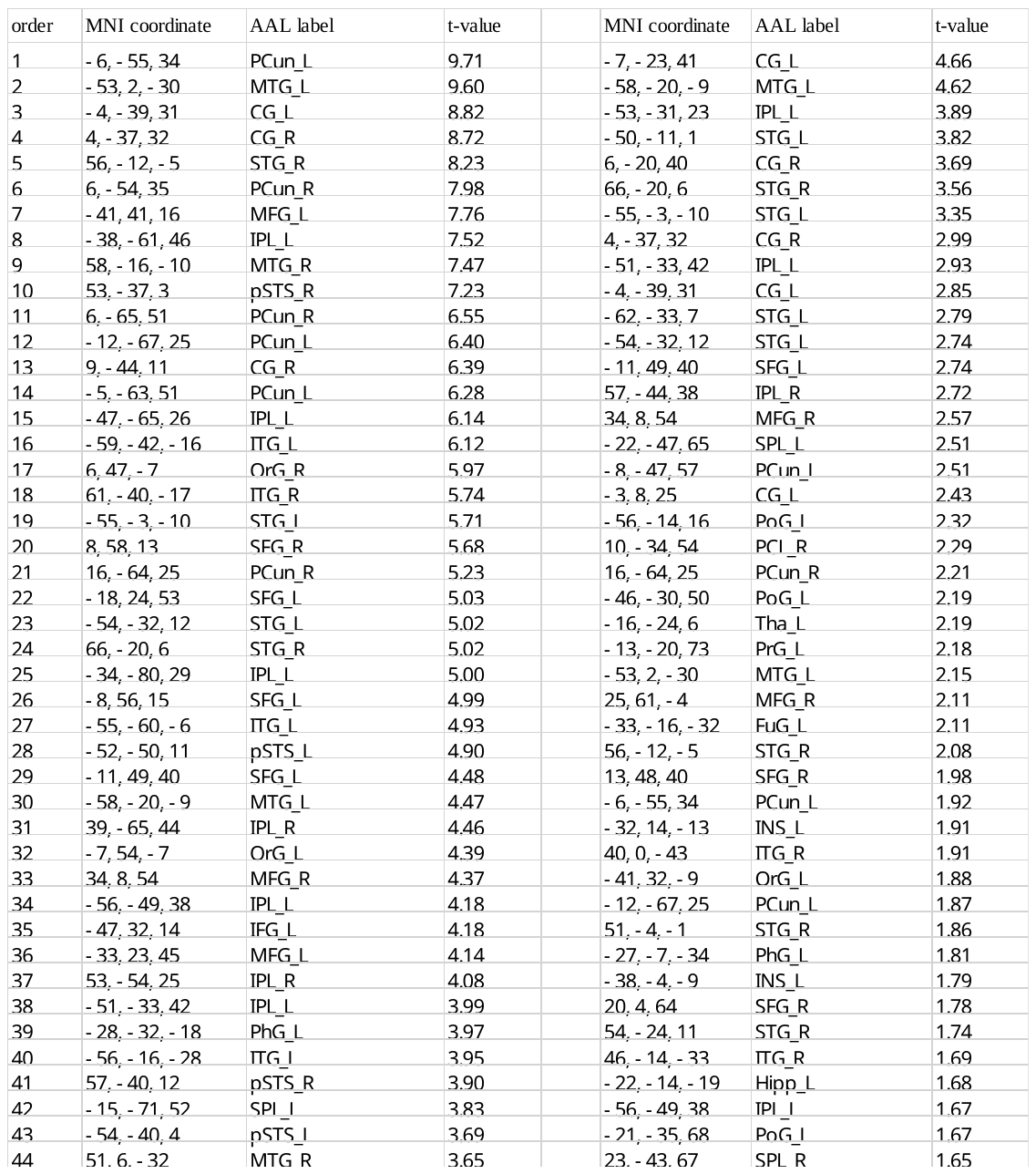

*Note.* SFG: superior frontal gyrus; MFG: middle frontal gyrus; IFG: inferior frontal gyrus; OrG: orbital gyrus; PrG: precentral gyrus; PCL: paracentral lobule; STG: superior temporal gyrus; MTG: middle temporal gyrus; ITG: inferior temporal gyrus; Fug: fusiform gyrus; PhG: parahippocampal gyrus; pSTS: posterior superior temporal sulcus; SPL: superior parietal lobule; IPL: inferior parietal lobule; PCun: precuneus; Pos: postcentral gyrus; INS: insular gyrus; CG: cingulate gyrus.
